# High Frequency of Nitroimidazole-Resistant *Trichomonas gallinae* in Competition Pigeons: Risk Factors and Therapeutic Implications

**DOI:** 10.64898/2026.01.30.702758

**Authors:** M. García-Piqueras, R. Suarez Lombao, P. Pérez Moreno, M. Bailén, D. Liebhart, M. González Clarí, M.T. Gómez-Muñoz, J. Sansano-Maestre

**Author notes:** Correspondence: María Teresa Gómez-Muñoz, Department of Animal Health, Faculty of Veterinary Sciences, University Complutense of Madrid, Avda. Puerta de Hierro s/n, 28040 Madrid, Spain., no Fax number available.

## Abstract

*Trichomonas gallinae* is a protozoan parasite of major concern in avian medicine, particularly in domestic pigeons (*Columba livia*). This study investigated the risk factors associated with the frequency of nitroimidazole resistance and *T. gallinae* prevalence in domesticated pigeons from Eastern Spain, kept for different competitions. A total of 220 pigeons from 11 lofts were sampled and examined by microscopy and culture, revealing a 63.6% infection prevalence. Genotyping identified genotype C as predominant, with occasional detection of genotype A, mixed A/C infections, and one isolate of Lineage III. *In vitro* susceptibility testing of 42 isolates showed a high prevalence (81%) of metronidazole resistance (MIC values ≥ 20 µg/ml), with minimum inhibitory concentrations (MICs) ranging from 5 to >100 µg/mL in 9/11 pigeon lofts examined. Resistance was significantly associated with the use of metronidazole and was more frequent in young and non-reproductive birds. Biannual treatments and the combination of ronidazole and dimetridazole at higher doses were associated with lower infection rates than monotherapies or annual treatments. No significant associations were found between resistance and environmental or loft management parameters, although poor hygiene and high bird density were common in lofts with resistant strains. These findings highlight the urgent need for regulated treatment protocols, improved biosecurity, and the development of alternative trichomonacidal agents to combat the emergence of drug-resistant *T. gallinae* in pigeon populations.

## 1. Introduction

The flagellated protozoan *Trichomonas gallinae* affects both wild and captive birds, leading to the disease known as avian trichomonosis (Quillfeldt et al., 2018). Columbidae family members are frequently infected (Marx et al., 2017), highlighting the domestic pigeon (*Columba livia*) as the main reservoir (Forrester & Foster, 2008). This parasite affects birds globally, including Eurasian collared dove (*Streptopelia decaocto*), stock dove (*Columba oenas*), European turtle dove (*Streptopelia turtur*), and wood pigeon (*Columba palumbus*), as relevant examples (Forrester & Foster, 2008; Lennon et al., 2013; Villanúa et al., 2006; Hegeman et al., 2007). Additionally, *T. gallinae* can infect other avian groups, including raptors and various passerine groups (Lawson et al., 2012). Direct disease transmission by different behavioural habits (adults feeding their young, courtship, and predation by birds of prey), or through the uptake of contaminated feed or water has been described. The oral route of infection during nutrition was identified as most common among adults (Lawson et al., 2012, Cole 1999).

Several parasite genotypes of oral trichomonads in birds have been described since Gerhold et al., (2008) published the first genetic classification in the USA, with two of the described genotypes predominant in Spain (Sansano-Maestre et al., 2009). Genotype C (according to Gerhold et al., 2008) was more frequent in Columbiformes, while genotype A was associated with higher virulence (Sansano-Maestre et al., 2009) and higher frequency in birds of prey (Martínez-Herrero et al., 2014, 2020). Additionally, infections produced by more than one genotype in the same animal have been reported (Zimre-Grabensteiner et al., 2010; Martínez-Herrero et al., 2019).

Infection prevalence varies significantly depending on the avian species and geographical location, reaching up to 74% in some European studies, with particular emphasis in pigeons from Germany, Spain, Malta, and Italy (Marx et al., 2017). Prevalence is also influenced by the diagnosis method employed, as the disease can be confirmed by oropharyngeal swabs observed under the microscope, cultured in TYM medium or amplified by PCR (Martínez-Herrero et al., 2021). In the 20^th^ century, nitroimidazoles were extensively used in sport pigeons, leading to drug resistance of the parasite (Franssen & Lumeij 1992; Munoz et al., 1998). While bacterial drug-resistance mechanisms are well documented, comparable studies in protozoa remain scarce (Santos et al., 2020, Caneschi *et al*., 2023). Nitroimidazole resistance threshold has been proposed for *T. vaginalis*, varying between aerobic or anaerobic conditions (12.5 in anaerobic *vs*. 38.8 µg/mL in aerobic conditions) (Narcisi & Secor,1996; Augostini et al., 2023). Since no established limit for *T. gallinae* has been established, this limits have been employed in previous *T. gallinae*-studies (Rouffaer et al., 2014), although Munoz et al reported a similar MIC value employing aerobic and anaerobic conditions in *T. gallinae* resistant isolates (Munoz et al, 1998).

Studies in recent years carried out on *Trichomonas gallinae* confirmed the presence of resistance around the world (Rouffaer et al., 2014; Cai et al., 2024).

In Spain, the impact of this parasite has been largely studied in Eastern regions due to the relevance of pigeon breeding for different sport events. Avian trichomonosis is well-recognised by the breeders, and most of them perform preventive or metaphylactic measures against the parasite mostly without veterinary supervision using different drug combinations according to their experience or the advice of other breeders.

In this context, the aim of the present study was to investigate the presence of *T. gallinae* and the frequency of nitroimidazole-resistant isolates related to the different pigeon breeding schemes, with a focus set on the evaluation of varying management practices, such as housing, treatment protocols, and individual parameters.

## 2. Materials and methods

### 2.1. Sampling: animals and pigeon lofts

The study area included eleven pigeon lofts, four from Valencia province, six from Alicante province, and one Castellón province, all within the Valencian Community in Spain. Letters were previously sent to the owners requesting their collaboration, and all who agreed to it were included in the present work. In this area there are different breeds of pigeons since there are two types of competition: one based on the morphology of the animals (fancy pigeons) and the other, called “Pica”, in which a group of trained males chase a female, guiding her to their pigeon loft (sporting pigeons).

A survey was conducted on the day of sampling to collect information on management measures, bird variables, and administered treatments. Management variables included the number of animals maintained in the loft (≤100 or >100 birds), floor type (fix or removable) and cleaning and disinfection frequency (weekly or monthly). From each bird recorded data included age, sex, reproductive status, whether it was in competition at sampling, and breed (morphology or Pica competitions). In cases where the birds were chemotherapeutically treated, the frequency (annual or biannual) of treatments and the drug combination were recorded. To prevent parasitosis, including trichomonosis, breeders routinely use metronidazole (alone or combined with amprolium or mebendazole) and ronidazole alone or combined with amprolium or dimetridazole, in this last case the manufacturer recommended dose of nitroimidazoles was double.

### 2.2. Culture and preservation of isolates

Sample collection was performed using oropharyngeal and crop samples collected with sterile swabs during clinical examination before inspected microscopically and cultured in TYM medium (Martínez-Herrero et al., 2017). The composition of this medium was as follows: 20 g trypticase (Sigma-Aldrich, St. Louis, MO, USA), 10 g D (+)-maltose (Sigma-Aldrich), 10 g yeast extract (Sigma-Aldrich), 1 g L-cysteine (Sigma-Aldrich), 0.1 g ascorbic acid (Sigma-Aldrich) and 10% inactivated fetal bovine serum (Sigma-Aldrich) per liter. After mixing all the ingredients, the pH was adjusted to 6.5 and sterilized by filtration through a 0.22 μm filter (Millipore, Billerica, MA, USA). Antibiotics and antimycotic supplements were used to avoid contamination: 24 ml nystatin/l (10,000 IU/ml, Sigma-Aldrich), 0.8 ml enrofloxacin/l (25 mg/ml, Bayer) and 5 ml/l chloramphenicol (100 μg/ml, Sigma-Aldrich) were added. The cultures were incubated at 37°C and daily reviewed under an inverted microscope to observe mobile trophozoites for up to four days. The cultures were passaged every 48 hours and after up to 5 serial passages, the isolates with living trichomonads were preserved in 5% dimethyl sulfoxide (DMSO) at −80°C and thawed for subsequent analyses as described below.

### 2.3. DNA extraction

After counting the parasites in a Neubauer chamber in triplicate, a total of 10^6^ trophozoites from each isolate were centrifuged to discard the culture medium and washed twice with PBS (600x*g*, 5 min each step). The pellet was employed to obtain DNA using the NZYTissue gDNA isolation kit (NZYTech, Lisbon, Portugal) according to the manufacturer’s protocol. The concentration of the obtained DNA was measured on a Multiskan GO microplate spectrophotometer with µDrop™ plates (Thermo Fisher Scientific, MA, USA) and stored at −20°C until use.

### 2.4. PCR of the fragment ITS1/5.8S/ITS2 and sequence analysis

Primers TFR1 (5′- TGCTTCAGCTCAGCGGGTCTTCC-3′) and TFR2 (5′- CGGTAGGTGAACCTGCCGTTGG-3′) were employed for PCR reactions to amplify a fragment of the ITS1/5.8S/ITS2 (Felleisen, 1997). PCR reactions were performed in 25 µL using 12.5 µL of Supreme NZYTaq II 2x Master Mix (NZYTech, Lisbon, Portugal). An initial step of 95°C for 5 min to activate the enzyme followed by 40 cycles of denaturation at 94°C for 30 sec, annealing at 66°C for 30 sec, extension at 72°C for 30 sec, and a final elongation step of 72°C for 10 min to allow the elongation of the PCR products were used. The results of PCR amplifications were visualized under UV light on 1% agarose gels stained with SYBR™ safe DNA gel staining (Invitrogen, Thermo Fisher Scientific, MA, USA).

The amplicons of the expected size were sent to Macrogen Spain (Madrid, Spain) for Sanger sequencing in forward and reverse directions. The resulting sequences were aligned using Molecular Evolutionary Genetics Analysis v.11 software (MEGA XI) (Tamura et al., 2021) and FinchTV 1.4.0 software (Geospiza, Inc.; Seattle, WA, USA), manually checked, and subjected to BLAST analysis using the nucleotide Basic Local Alignment Search Tool (BLAST; National Library of Medicine, Rockville, MD, USA).

Only clear sequences with more than 150 bp in both directions were sent to GenBank for accession numbers.

### 2.5. *In vitro* evaluation of metronidazole resistance

Evaluation of the metabolic activity of trophozoites was performed using the MTT (3-[4,5-dimethylthiazol-2-yl]-2,5 diphenyl tetrazolium bromide) method in 96-well plates with round bottom, following the procedure previously described (Bailén *et al*., 2022). Briefly, the assay was performed in quadruplicate, after counting the trophozoites as described above, a total of 150 µl/per well of a solution containing 5×10^5^ *T. gallinae* trophozoites/ml in TYM medium was added to the plate. Routinely, metronidazole was added to each well at different final concentrations, from 40 to 0.1 µg/ml. In cases of high resistance to metronidazole, concentrations up to 100 µg/ml were also evaluated. As controls, medium and culture of each isolate without metronidazole (Acros Organics, Madrid Spain) were employed. The plate was then sealed with a sterile waterproof and air-tight adhesive lid and incubated at 37°C for 24 h. Previous assays comparing this procedure with Vaseline sealed wells under our test conditions offered the same MIC values, facilitating MTT assays and MIC determination at the same time. After sealing and incubation, the plate was centrifuged at 750 x *g* to eliminate the medium. One hundred µl of a solution containing MTT (1.25 µg/ml) (Acros Organics, Madrid Spain) and PMS (phenazine methosulfate at 0.1 µg/ml, Acros Organics, Madrid Spain) was added to each well and incubated at 37°C to allow formazan salts to form. After 30 minutes, 50 µl of a HCL-SDS solution (37 µl of 1N HCL in 100 ml of 10% sodium dodecyl sulphate) was added to each well to dissolve formazan salts. After thirty minutes of incubation at 37°C, the plate was read in a spectrophotometer at 570 nm (Multiskan GO microplate spectrophotometer, (Thermo Fisher Scientific, MA, USA). Then, the following formula was applied to calculate the percentage of activity of the drug:

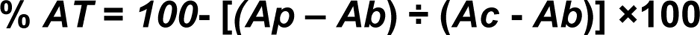

Ap is the absorbance of the tested well, Ab is the absorbance of the blank well, and Ac is the absorbance of the control well (each isolate culture without drug).

### 2.6. Determination of MIC value and IC_50_ value

Two assays were performed for each isolate in quadruplicate to determine MIC and IC_50_ values. The minimum inhibitory concentration (MIC) was established at the lowest concentration of the drug that prevents visible movement (a dose immediately higher to the highest dose that shows trophozoite movement). For that purpose, the plates from the MTT assay were observed in an inverted microscope before the addition of MTT solution at 24 h of exposure to metronidazole at the above-mentioned concentrations. Since there is no standardized threshold to establish metronidazole resistance in *T. gallinae*, limits for *T. vaginalis* were employed. MIC values >12.5 µg/mL established for anaerobic conditions (Narcisi & Secor,1996) and employed in previous *T. gallinae*-studies (Rouffaer et al., 2014), were used at threshold, Inhibitory concentration 50 (IC_50_) was determined as the concentration of metronidazole able to inhibit 50% of the metabolic activity of each isolate. A linear regression analysis (% cell viability on log dose) was calculated with the absorbance values of each drug concentration employing STATGRAPHICS Centurion XIX (https://www.statgraphics.com).

### 2.7. Statistical Analysis

Categorical variables were described using absolute and relative frequencies. Mean and standard deviation (SD) were expressed as continuous variables. Variables were classified according to their relationship with management practices in the loft (loft size, loft modality, floor type, cleaning and disinfection frequency). Age, sex, reproductive status, breed, and whether they were competing at the time of sampling were recorded as intrinsic variables from animals. Finally, variables related to metaphylactic measures recorded included the administration or non-administration of antiparasitic drugs, the frequency of treatments, and the administered drug combination.

Associations between categorical variables were explored using chi-square tests and Cramer’s V to describe the strength of association. To assess multicollinearity among candidate predictors, variance inflation factors (VIF) were computed on the design matrix with appropriate dummy coding; tolerance < 0.20 or VIF > 5 indicated collinearity and such predictors were excluded or combined. The associations between *Trichomonas* infection and predictors were analysed using a Generalised Estimating Equations (GEE) model (logit link), using the pigeon loft as a clustering factor, assuming an exchangeable working correlation structure. Odds ratios (OR) and their 95% confidence intervals (95% CI) were calculated. P-values were obtained using robust variance. For the resistance analysis, risk ratios (RR) were calculated to provide a descriptive measure of the strength and direction of association between exposure variables and the presence of resistance. These RR values were interpreted as descriptive rather than inferential since the study design was cross-sectional and not a cohort study. Statistical significance was set at two-sided p < 0.05. All statistical analyses were performed with SPSS 20.0. (IBM Corp., Armonk, NY, USA).

### 2.8. Ethics Committee approval

Samples were collected under the authorization of the ethics committee approval number PROEX 166.7/21.

## 3. Results

### 3.1. Prevalence of *T. gallinae* and analysis of risk factors

A total of 220 animals from 11 different lofts were included in the study (Table 1). Most of the lofts were classified as small size (≤100 animals) (68.2%), with removable floor enclosures (59.6%) and with weekly cleaning (56.4%). The drinking water source in all pigeon lofts was cleaned or changed without a regular schedule and for this reason this parameter was not included in the analysis. Regarding bird parameters, most of them were males (61.8%), adults (67.7%), not in reproduction (59.1%), bred for Pica competitions (82.3%), and most of them were in competition at sampling (95%). Concerning therapeutic measures, only two lofts (11.8% of birds, n=26) reported not using nitroimidazoles against *T. gallinae*. The combinations of drugs employed by the nine pigeon breeders who treated their animals (n=194) were: exclusively metronidazole (25.8% of treated birds, n=50, one pigeon loft), metronidazole combined with amprolium and mebendazole (21.7% of treated birds, n=42, two pigeon lofts), ronidazole + dimetridazole (22.7% of treated birds, n=44, two pigeon lofts), and ronidazole + amprolium (29.9% of treated birds, n=58, four pigeon lofts). The frequency of medication administration also varied between facilities that medicate birds, using one (37.6%, n=73, four pigeon lofts) or more than one treatment per year (62.4%, n=121, eight pigeon lofts), with most of them applying biannual treatments (60.3%, n=117, 7 pigeon lofts). In general, a high variation in preventive measures and administered treatments was observed, without veterinarian supervision.

**Table 1.**
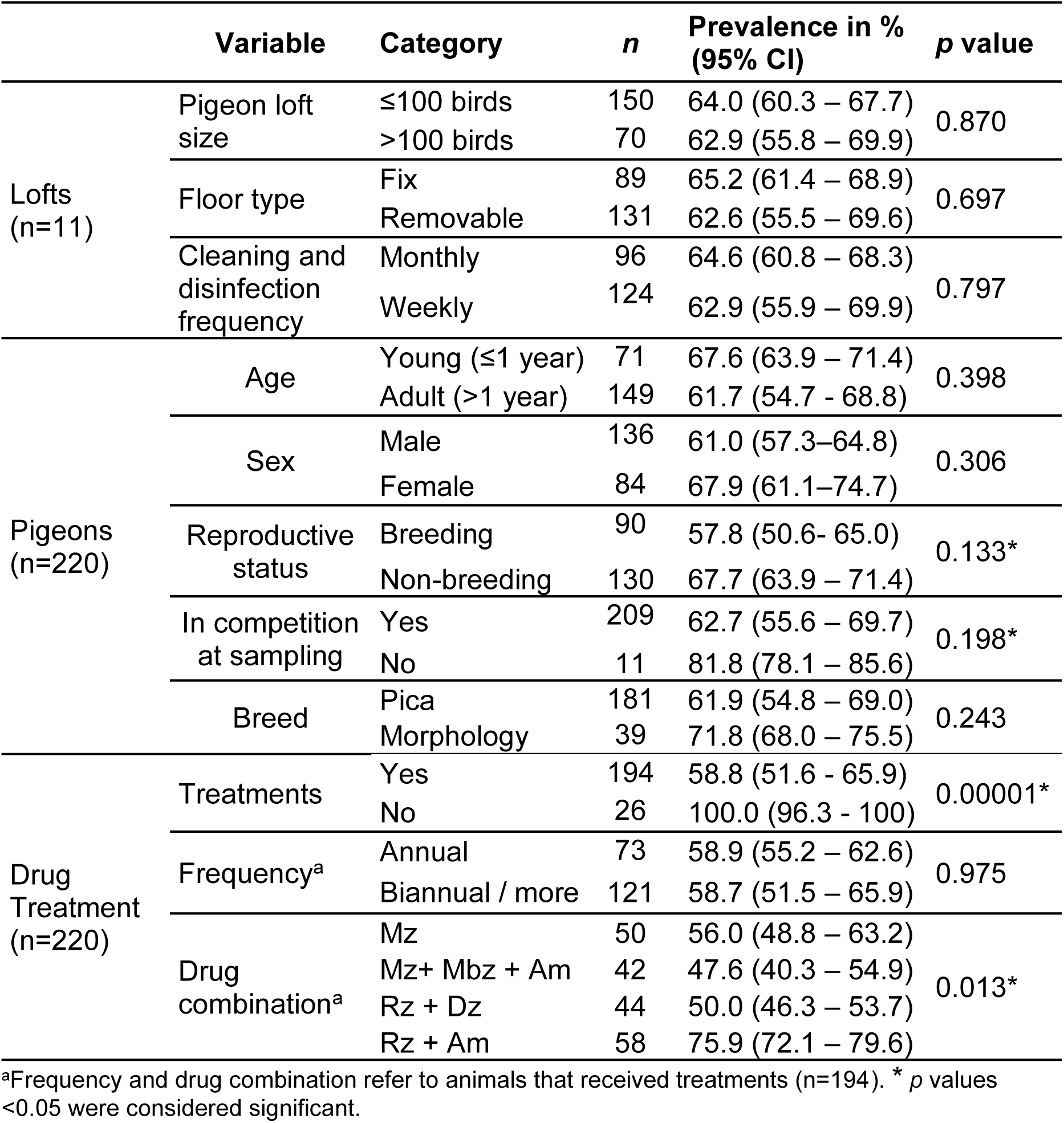
Categorized qualitative variables, number of pigeons sampled and influence on *T. gallinae* prevalence. Significance of association between each variable and *T. gallinae* prevalence is indicated in the last column (*p* values). GEE model included variables with *p* value < 0.2 or biologically relevant (*). Mz, metronidazole; Dz, dimetridazole; Rz, ronidazole; Am, amprolium; Mbz, mebendazole.

Overall, 140 of 220 swabs from the upper digestive tract were positive by direct microscopic examination, yielding in a prevalence of 63.6%, but some of the positive samples could not be maintained in culture due to poor growth or bacterial or fungal contamination. A total of 42 isolates from 9 pigeon lofts were preserved from these samples for further analysis.

*T. gallinae* was detected in all pigeon lofts, although prevalence values varied depending on the facility, ranging from 42.9% to 100% of the sampled birds. No significant correlation was identified between *T. gallinae* infection and the evaluated management variables. Hence, the size of the facility (64.0% *vs*. 62.9%), the type of floor (65.2% *vs.* 62.6%), and the frequency of cleaning (64.6% vs. 62.9%) did not significantly influence the parasite prevalence (Table 1). Similarly, *T. gallinae* prevalence was not significantly altered by other variables analyzed (Table 1). Pigeons participating in morphology competitions had a prevalence of 71.8%, while animals that did not participate in competitions at the time of sampling showed a higher prevalence, 81.8%, although the small sample size in the latter group limited statistical comparison (n=11, *p*= 0.198). Young pigeons presented a higher prevalence (67.6%) compared to adults (61.7%), but the difference was non-significant (*p* = 0.398). Females had a higher infection rate (67.9%) than males (61.0%), without statistical significance (*p*= 0.306). Similarly, non-reproductive pigeons showed a slightly higher prevalence (67.7%) compared to reproductive individuals (57.8%), with no significant association (*p*= 0.133).

However, a significant association with the prevalence of the parasite in the lofts was drug administration (p< 0.001). All animals analysed from lofts that did not use anti-*Trichomonas* treatments (n=26) were highly positive for *T. gallinae* infection (prevalence value= 100%) compared to pigeons from treated lofts (prevalence value= 58.8%). Therefore, the absence of treatment was associated with a higher risk of infection, although it was not possible to estimate odds ratios as there was no variation in the outcome.

All nitroimidazole treatments were administered via drinking water. Among the 194 pigeons that received antiparasitic treatment, 114 tested positives for *T. gallinae* by microscope observation. For these treated pigeons, the GEE model included variables with tolerance values above 0.20, VIF values below 5 and p ≤ 0.2 (reproductive status, competition, and drug combination) (Table 2). Frequency of drug administration (once or more per year), despite not being statistically significant, was included for its relevance in the efficacy of the treatment. As a result, the model combining two nitroimidazoles (ronidazole and dimetridazole) was associated with lower *T. gallinae* infection rates (OR= 0.77, 95% CI= 0.67-0.89), whereas pigeons treated with only one nitroimidazole (ronidazole combined with amprolium) were twice as likely to be infected as birds treated with metronidazole (OR= 2.13; 95% CI= 1.11-4.17). The combination of metronidazole with other non-nitroimidazole drugs (mebendazole and amprolium) did not improve the efficacy of metronidazole alone against the parasite (OR= 0.79; 95% CI= 0.32-1.92). The administration of two treatments per year was considered a potential protective factor, after eliminating confounding factors in the GEE model (Table 2), being one treatment per year a potential risk factor (OR= 1.32; 95% CI= 1.05-1.64).

**Table 2.**
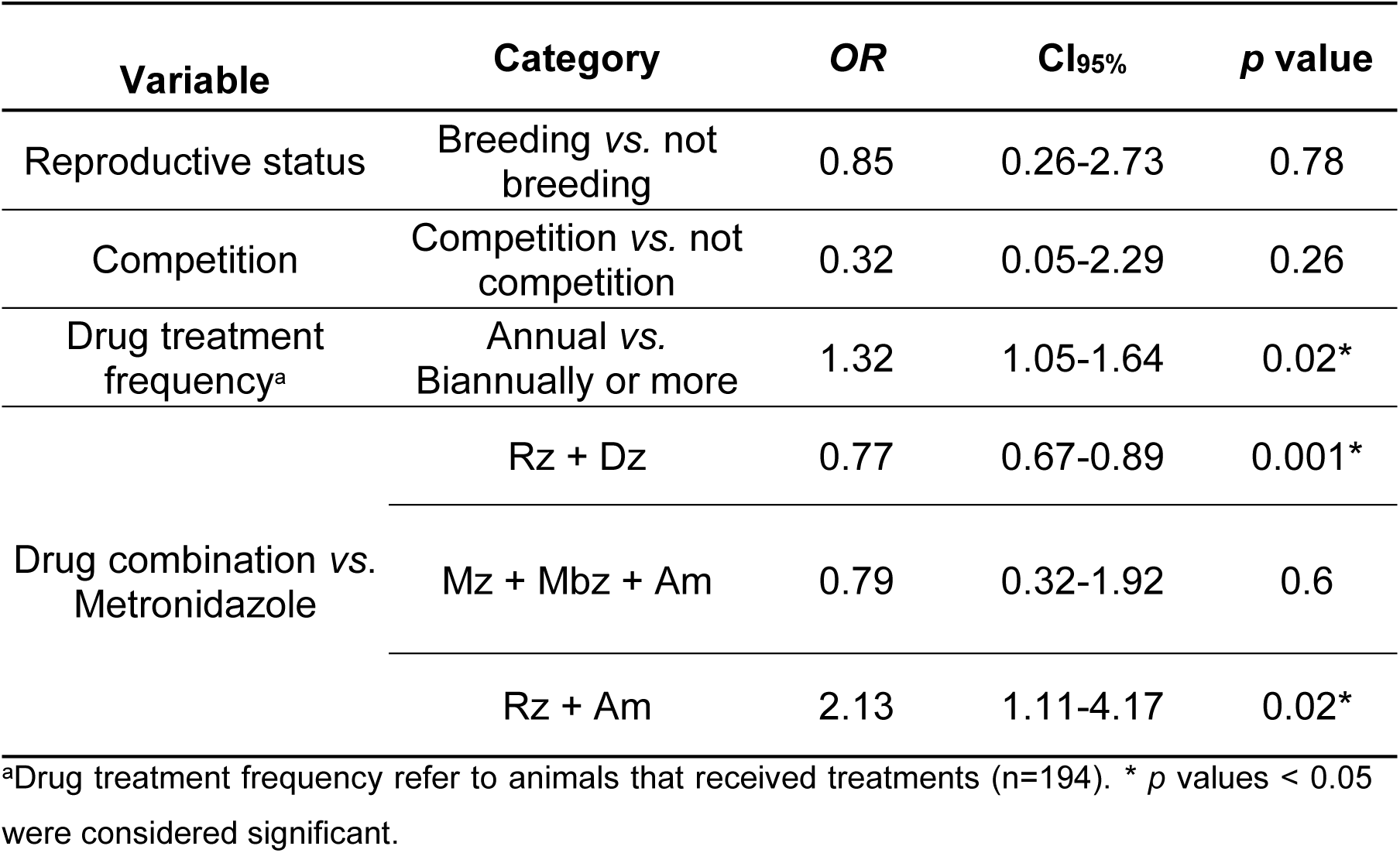
Association of selected variables on *T. gallinae* prevalence employing a GEE model.

### 3.2. Genotyping and GenBank accession numbers

Most of the obtained isolates displayed genotype C (n=25), according to Gerhold et al. (2018) and only two isolates displayed genotype A (isolates 276 and 277) (Table 3). A minor proportion displayed a mixed genotype (A/C, n=3) and were not sent to GenBank. Sequence from isolate 228 displayed 100% coverage and 100% identity with sequence KX459507.1, isolated from a European turtle dove (*S. turtur*) and named as lineage III previously (Marx et al., 2017). GenBank accession numbers obtained in this study are PX936831-62.

**Table 3.** Genotypes, IC_50_ values (µg/ml), MIC values (µg/ml), and pigeon loft number of the analyzed isolates. Nitroimidazole-resistant isolates (MIC values >12.5 µg/ml) are indicated with “Yes” whereas non-resistant isolates are indicated with “No”.

| Isolate (pigeon loft number) | Genotype | IC<sub>50</sub> (µg/ml) (95 % CI) | MIC (µg/ml) | Resistance |
| --- | --- | --- | --- | --- |
| Val-23-1 (11) | N.d.<sup>a</sup> | 4.3 (3.6 -5.3) | 20 | Yes |
| Val-23-2 (11) | Mixed (A/C) | 10.8 (9-13) | >40 | Yes |
| Val-23-3 (11) | Mixed (A/C) | 6.1 (5.4 -6.9) | 40 | Yes |
| Val-23-4 (11) | C | 1.4 (1.0-2) | 5 | No |
| Val-23-7 (11) | C | 1.3 (1.0-1.6) | 10 | No |
| Val-23-9 (11) | N.d.<sup>a</sup> | 0.5 (0.8- 1.0) | 10 | No |
| Val-23-10 (11) | C | 2.9 (2.2 – 3.8) | >40 | Yes |
| Val-23-12 (11) | N.d.<sup>a</sup> | 11.5 (8.7-15.3) | 10 | No |
| Val-23-25 (10) | C | 0.6 (0.2 -1.5) | 40 | Yes |
| Val-23-40 (7) | C | 0.6 (0.4- 0.9) | 10 | No |
| Val-23-46 (7) | N.d.<sup>a</sup> | 2.5 (1.6 – 3.9) | 40 | Yes |
| Al-23-88 (3) | N.d.<sup>a</sup> | 7.6 (5.1 -11.4) | >40 | Yes |
| Al-23-90 (2) | C | 26.9 (21.9 -33) | >100 | Yes |
| Al-23-93 (2) | C | 4.7 (3.1 -6.9) | >40 | Yes |
| Al-23-105 (2) | C | 5.4 (4.3 – 6.9) | >40 | Yes |
| Al-23-169 (1) | Mixed (A/C) | 1.9 (1.3 -2.8) | >40 | Yes |
| Val-23-195 (11) | C | 3.8 (2.5 – 5.8) | >40 | Yes |
| Val-23-199 (11) | C | 3.4 (2.2 -5.2) | 10 | No |
| Val-23-207 (11) | C | 1.3 (0.9- 2) | 10 | No |
| Val-23-208 (11) | N.d.<sup>a</sup> | 3.8 (2.8 -5) | 10 | No |
| Val-23-210 (11) | C | 3.6 (2.8- 4.5) | 20 | Yes |
| Val-23-211 (11) | C | 2.1 (1.4 -3.1) | 20 | Yes |
| Val-23-212 (11) | C | 1.4 (1.1 – 2) | 40 | Yes |
| Val-23-215 (11) | C | 4.7 (3.5 – 6.4) | 20 | Yes |
| Val-23-217 (11) | C | 0.7 (0.3 – 1.3) | 20 | Yes |
| Val-23-218 (11) | C | 7.7 (5.5 – 10.7) | 40 | Yes |
| Val-23-220 (11) | C | 7.4 (6.2 – 8.8) | >40 | Yes |
| Val-24-228 (5) | Lineage III | 54.3 (37.8 – 78.0) | >80 | Yes |
| Val-24-229 (5) | C | 9.3 (6.9 – 12.7) | 40 | Yes |
| Val-24-244 (5) | C | 24.7 (16.5- 37.1) | 80 | Yes |
| Val-24-247 (5) | C | 3.6 (3.0-4.3) | 20 | Yes |
| Al-24-261 (9) | C | 82.6 (52.4 – 130.1) | >80 | Yes |
| Al-24-264 (9) | C | 48.7 (34.1. -69.7) | >80 | Yes |
| Al-24-272 (9) | C | 5.8 (4.0-8.6) | 50 | Yes |
| Al-24-273 (9) | C | 7.0 (6.0- 8.2) | 20 | Yes |
| AI-24-276 (9) | A | n.d. | >80 | Yes |
| AI-24-277 (9) | A | 49.7 (34.8 -52.5) | 80 | Yes |
| AI-24-279 (9) | N.d.<sup>a</sup> | 47.6 (19.8 - 114.8) | >80 | Yes |
| AI-24-281 (9) | C | 14.6 (10.1- 21-1) | >80 | Yes |
| Cas-24-292 (4) | C | n.d. | 80 | Yes |
| Cas-24-296 (4) | C | 4.4 (2.4 -8) | 50 | Yes |
| Cas-24-299 (4) | C | 11.9 (9.2 -15.3) | 100 | Yes |

<sup>a</sup>Not determined due to unsuccessful growth

### 3.3. Frequency of resistant isolates

A total of 42 *T. gallinae* isolates from nine lofts were evaluated for phenotypic resistance to metronidazole using both MIC and IC₅₀ values, although two isolates did not grow properly and IC_50_ values could not be estimated. IC₅₀ values were used as complementary indicators of reduced sensibility. MIC values ranged from 5 to >100 µg/mL, while IC₅₀ values ranged from 0.5 to 82.6 µg/mL, reflecting wide variability in susceptibility to metronidazole (Table 3). According to the resistance threshold, 34 isolates were classified as resistant, and eight were classified as susceptible (MIC ≤12.5 µg/mL). In general, a positive correlation between the MIC and IC_50_ values was observed (Spearman’s p = 0.667, p< 0.0001). Among the 34 resistant isolates from 9 pigeon lofts, 10 obtained from 3 pigeon lofts showed high values for both MIC (≥80 µg/ml) and IC₅₀ (11.9-82.6 µg/ml), while the remaining isolates were classified as resistant with MIC values from 20 to 50 µg/ml and IC_50_ values from 0.6 to 10.8 µg/ml. These findings confirm a high prevalence of resistance to metronidazole among the isolates analyzed.

Pigeon loft-related variables did not show any significant association with the presence of resistant isolates and were not included in the multivariable analysis, although most of resistant strains were recovered from lofts with poor hygiene (monthly cleaning) (82.4%) and high bird density (>100 birds) (70.6%). However, host-related variables age, reproductive status, and competition modality showed overall moderate associations with the frequency of resistant isolates (p<0.2). Drug treatments were regrouped according to the nitroimidazole present in their composition, resulting in 25 strains isolated from pigeons treated with ronidazole (59.5%) (one of them treated with ronidazole and dimetridazole), 12 strains from pigeons treated with metronidazole (28.6%) and five obtained from animals without treatment (11.9%). Among the treated pigeons (n=37), 11 received one dosage (29.7%) and 26 received two or more treatments throughout the year (70.3%) (Table 4). Associations between some of the selected variables and *T. gallinae* resistance were observed. Young pigeons were associated with resistant strains when compared with adults (RR = 1.28, 95% CI= 1.01–1.63). Similarly, non-breeding pigeons were associated with resistant strains in comparison with breeding birds (RR = 1.33; 95% CI= 1.01-1.76). Competition modality did not show a significant association with nitroimidazole resistance of the strains, although birds belonging to the Pica competition modality tend to be associated with higher frequency of nitroimidazole resistance.

**Table 4.** Relationships between selected qualitative variables and presence of resistant isolates (MIC value > 12.5 µg/mL). RR higher than 1, including 95% CI, were considered risk factors.

| Variable | Category | n | RR (95 % CI) | p value | Risk factor |
| --- | --- | --- | --- | --- | --- |
| Age | Young (<1 year) | 10 | 1.28 (1.01 – 1.63) | 0.039* | young |
|  | Adult (>1 year) | 32 |  |  |  |
| Reproductive status | Reproductive | 24 | 1.33 (1.01 – 1.76) | 0.04* | non-reproductive |
|  | Non-reproductive | 18 |  |  |  |
| Competition modality | Pica | 34 | 1.74 (0.87 – 3.56) | 0.11 | Pica (risk tendency) |
|  | Morphology | 8 |  |  |  |
| Treatment frequency<sup>a</sup> | Biannual | 26 | 1.39 (0.87 – 2.22) | 0.167 | biannual (risk tendency) |
|  | Annual | 11 |  |  |  |
| Drug<sup>a</sup> | Metronidazole | 25 | 1.39 (1.09 – 1.77) | <0.001* | metronidazole |
|  | Ronidazole | 12 |  |  |  |

<sup>a</sup>Drug and treatment frequency were assessed only in treated animals (n=37). \* p values < 0.05 were considered significant.

Regarding treatment frequency, there was no statistically significant association with the frequency of metronidazole-resistant strains; however, point estimates suggested a positive trend toward an association between resistance and more administrations/year (RR = 1.39, 95% CI= 0.87 −2.22). A consistent and statistically significant association was found for the nitroimidazole used: pigeons treated with metronidazole had 1.39-fold higher risk of developing metronidazole resistance compared with those treated with ronidazole (RR = 1.39, 95% CI= 1.09 - 1.77).

## 4. Discussion

The overall prevalence of *T. gallinae* infection in this study is 63.6%, confirming its widespread presence among domestic pigeons in eastern Spain. This value aligns with prior findings in the Iberian Peninsula, although significant variation exists depending on region, bird population, and diagnostic methodology. In the same geographic area, Sansano-Maestre *et al*. (2009) reported a prevalence of 44.8**%** in pigeons by culture in TYM medium, although the sampled birds were not racing pigeons. More broadly, studies across Spain have shown varying prevalence depending on pigeon species and epidemiological context. As examples, Villanúa *et al*. (2006) reported 34.2% prevalence in wild wood pigeons, considering the presence of lesions together with parasite culture and microscope observation, while Marx *et al*. (2017) detected 70% prevalence in domestic pigeons across several European countries employing culture and PCR. While culture is particularly useful to obtain isolates, PCR offers a higher sensitivity to detect positives to *T. gallinae* infection (Martínez-Herrero et al., 2021). Also, the higher flock density of domestic sport pigeons is a risk factor for higher prevalence when compared to wild populations, a fact that has also been pointed out by Rouffaer et al (2014). Three of our isolates displayed a mixed genotype, which can be explained by the presence of genetically different clonal isolates present in the same bird (Zimre-Grabensteiner et al., 2011), a common finding as already described in previous prevalence studies (Sansano-Maestre et al. 2009; Alrefaei et al., 2019; Martínez-Herrero et al., 2021).

However, most of the previous studies have not assessed the role of routine treatment protocols or preventive management practices in shaping *T. gallinae* prevalence in domestic pigeons. In our study, management-related variables, including loft size, floor type, and cleaning frequency, were not significantly associated with infection risk. Similarly, intrinsic bird-level factors such as reproductive status and competition participation did not show significant associations. However, the limitation of the study is its cross-sectional design and the voluntary participation of pigeon breeders, hence, a potential selection bias must be considered. Notably, none of the surveyed lofts operated under veterinary supervision nor control drug dose intake, and none implemented regular cleaning of drinking water sources—factors that likely contribute to the persistence and transmission of *T. gallinae* within loft environments (Grabensteiner et al., 2010; Sansano-Maestre et al., 2009).

The only factor significantly influencing the prevalence of *T. gallinae* infection in lofts was the absence of antiparasitic treatment, since the infection reached 100% in untreated individuals, compared to 58.8% in treated birds (*p*<0.001). This confirms previous findings that highlight antiparasitic therapy as a key strategy for controlling *T. gallinae* transmission in pigeon lofts (Amin et al., 2014). However, the efficacy of administered treatments also varied according to the protocol and the frequency of administration. Thus, compared with metronidazole alone, lofts which employed ronidazole alone had higher odds to harbor animals with *T. gallinae* and had the highest parasite prevalence (38.6%). On the contrary, lofts treated with a combination of two nitroimidazoles (ronidazole + dimetridazole) displayed the lowest percentage of *T. gallinae* infected birds (19.3%) and this combination of nitroimidazoles was considered a potential protective factor. Treatments based on combinations of metronidazole and other non-nitroimidazole drugs, such as amprolium or mebendazole, did not show differences in their efficacy compared with metronidazole alone. These variations in efficacy could be due to the nitroimidazoles used (monotherapy or combined therapy). Given that one nitroimidazole alone (ronidazole alone or in combination with non-nitroimidazole drugs) was not associated with parasitism control, the combination of two nitroimidazole drugs (ronidazole+dimetridazole) appears to be associated with lower *T. gallinae* infection rate. This combination implied receiving a higher dose of nitroimidazoles and in cases of unresponsiveness to the regular doses of nitroimidazoles, the effectiveness of increased dosages of the drug has been reported (Franssen & Lumeij, 1992).

The efficacy of repeat treatments in controlling trichomonosis has been poorly explored. In our case, biannual preventive treatments were associated with lower infection rates compared to annual treatments, since pigeons from lofts with a single preventive treatment were associated with higher frequency of *T. gallinae* infections than pigeons treated twice, after eliminating confounding factors in the GEE model. Repeated treatments, however, have been linked to the appearance of nitroimidazole resistance (Lumeij and Zwijneneberg, 1990; Rouffaer et al., 2014; Munoz et al., 1998; Duchatel & Vindevogel, 1998; Gómez-Muñoz et al., 2022). In most cases of drug resistance, a recommendation of drug rotation has been made to control and reduce the selection pressure for resistance. However, it must be considered that most of the nitroimidazole-resistant isolates (up to 85% of resistant isolates in some studies) are resistant to several nitroimidazoles, such as metronidazole, dimetridazole, ornidazole, carnidazole, or ronidazole (Munoz et al., 1998, Franssen & Lumeij 1992, Rouffaer et al., 2014, Cai et al., 2024), and no alternatives exist to treat avian trichomonosis.

In the present study, resistant isolates (MIC values of 20 µg/ml or higher in our conditions) were recovered in 9/11 lofts, mostly from treated birds with nitroimidazoles, either alone or in combination with other drugs such as amprolium and mebendazole. However, four of five strains from untreated animals from two lofts were also resistant to metronidazole. This suggests that resistant strains are widespread among the lofts. The origin of resistance could be treatment failures, subtherapeutic dosing, or selection of resistant subpopulations following repeated or poorly regulated drug exposure, as suggested in previous studies (Lumeij & Zwijnenberg, 1990). The results from the present survey suggest that all breeders have conducted preventive treatments at some point, although some of them stopped doing so when they did not observe improvements in the performance of their animals. This could have led to the selection of persistent resistant strains in pigeon lofts. A common fact is that all breeders administered preventive treatments using drinking water. This route is easy to use when treating multiple animals, but it does not allow for verification of the amount of medication ingested, so underdosing is common, favoring drug resistance (Gómez-Muñoz et al., 2022). A widespread problem that was observed is a therapy switch without veterinary supervision. The mentioned drugs can be obtained by online purchase, although nitroimidazole drugs are subject to veterinarian supervision in Spain as well as in the EU and forbidden for preventive treatment (Regulation EU 2019/6, 2019). Our study showed that young animals harboured resistant strains more frequently than adults, in agreement with observations of Cai et al (2024), who reported resistant strains more frequently in young pigeons (<30 days old).

We observed that non-reproductive animals carried resistant strains more frequently than breeding pigeons. Chronic stress, produced by captivity and competitions, may produce significant detrimental effects to individuals, such as increasing the susceptibility, leading to higher parasite loads and infection that could predispose these animals to concentrate these circulant strains, perpetuating the transmission cycle within the lofts (Nordling et al. 1998).

Risk analysis did not show significant relationships in the appearance of resistance with the frequency of treatment administration, despite a positive trend association being observed in animals receiving two treatments per year and nitroimidazole resistance. Regarding the drugs employed, metronidazole was associated with a 1.39-fold higher risk of resistance compared to ronidazole. This drug is by far the most widely used by breeders because it was the most easily accessible in Spain. Anyhow, in recent years, due to online trade, it has been possible to access other nitroimidazoles formulated for pigeons without veterinary supervision. This scenario needs to change in the context of the EU legislation (Regulation EU 2019/6).

Molecular genotyping revealed that genotype C predominated among resistant isolates, but also in non-resistant isolates. In Spain, genotype C predominates in domestic pigeons compared with wild birds in previous studies (Sansano-Maestre et al., 2009, Martínez-Herrero et al., 2014). However, two highly resistant isolates belonged to genotype A (e.g., isolates 276, 277), one to Lineage III (isolate 228), and three mixed genotypes (A/C) were identified in pigeons from pigeon lofts #1 and #11. This diversity suggests that resistance is not confined to a specific genotype, in contrast to the suggestion made by Cai *et a*l (2024), who found more resistance associated with genotype A. Our findings agree with the observations made by Rouffaer et al. (2014), who found more frequently resistant isolates with genotype C, which was also the most frequent genotype isolated from pigeons.

Among the eight *T. gallinae* isolates classified as susceptible to metronidazole (MIC ≤ 12.5 µg/mL), genotyping data were available for six of them. All were identified as genotype C, which is also frequent in resistant isolates.

Environmental and husbandry factors were analyzed as factors influencing the resistance dynamics, but no significant results were obtained. Most resistant isolates were recovered from pigeons housed in lofts with monthly cleaning, solid uralite flooring, and high population densities. Notably, in pigeon loft 9, which yielded resistant isolates with the highest IC₅₀ values, exhibited all these conditions. These findings point to the cumulative impact of suboptimal hygiene, crowding, and drug pressure in fostering resistance development and maintenance.

The growing evidence of nitroimidazole resistance among *T. gallinae* isolates calls for urgent implementation of alternative control strategies. While nitroimidazoles remain the cornerstone of trichomonosis treatment in birds, their efficacy is increasingly compromised, particularly in settings of chronic exposure, underdosing, poor biosecurity measures, and absence of veterinary supervision. Enhanced hygienic measures are essential for reducing transmission pressure within lofts. Frequent removal of feces, regular disinfection of drinkers and feeders, and minimizing overcrowding can significantly decrease environmental contamination and oral exposure to trophozoites (Amin et al., 2014).

Pharmacological alternatives should be considered. A study from Sheng Fan Jing *et al*. (2023) identified anisomycin, fumagillin, and MG132 (a synthetic peptide aldehyde) as promising drug candidates against *T. gallinae*. These compounds demonstrated strong *in vitro* inhibition with low toxicity. Preliminary studies have shown promising activity *in vitro* of certain essential oils (e.g., *Lavandula*, *Origanum*, *Thymus*, *Salvia*, and *Satureja* essential oils) (Bailen et al., 2022) and plant-derived extracts against *T. gallinae* (Malekifard *et al*., 2021). Other candidates under exploration include benzimidazole derivatives, silver nanoparticles, and ionophore compounds, which may provide trichomonacidal activity through non-nitroimidazole pathways (De Brum et al., 2017; Heshmati *et al*. 2023). However, these agents require rigorous *in vivo* testing to establish safety, efficacy, and pharmacokinetic profiles in avian hosts.

Based on our findings, we recommend a multimodal approach to control *T. gallinae* in domestic pigeons bred for sport, particularly in settings where similar treatment regimens are employed. This should include higher doses of antiparasitic drugs or combined nitroimidazole therapy, strict hygiene protocols—particularly regarding drinking water and loft sanitation—periodic susceptibility testing, enhanced veterinary oversight, and tighter regulation of over-the-counter access to nitroimidazoles. These strategies are essential to preserve the efficacy of existing treatments and to curb the spread of resistant *T. gallinae* strains within and beyond pigeon populations. Further research is necessary to evaluate alternative therapeutic compounds and to establish surveillance systems for monitoring resistance in avian trichomonosis.

## Authors contribution

Conceptualization and design: JSM and MTGM. Acquisition of data: MGP, RSL, PPM, JSM. Analysis and interpretation of data: MGP, MB, RSL, PPM, MGC, DL. Funding acquisition: MTGM, JSM. Draft of the article: MGP, JSM and MTGM. Critical revision for important intellectual content: JSM, DL, and MTG. Final approval: all the authors.

